# Multilocus phylogeny and historical biogeography of *Hypostomus* shed light on the processes of fish diversification in La Plata Basin

**DOI:** 10.1101/2020.09.22.308064

**Authors:** Yamila P. Cardoso, Luiz Jardim de Queiroz, Ilham A. Bahechar, Paula E. Posadas, Juan I. Montoya-Burgos

## Abstract

Distribution history of the widespread Neotropical genus *Hypostomus* to shed light on the processes that shaped species diversity. We inferred a calibrated phylogeny; ancestral habitat preference, ancestral areas distribution, and the history of dispersal and vicariance events of this genus. The phylogenetic and distributional analyses indicate that *Hypostomus* species inhabiting La Plata Basin do not form a monophyletic clade, suggesting that several unrelated ancestral species colonized this basin in the Miocene (∼17 Mya). Dispersal to other rivers of La Plata Basin started about 8 Mya, followed by habitat shifts and an increased rate of cladogenesis. Amazonian *Hypostomus* species colonized La Plata Basin several times in the Middle Miocene, probably via the Upper Paraná and the Paraguay rivers that acted as biogeographic corridors. During the Miocene, La Plata Basin experienced marine incursions; and geomorphological and climatic changes that reconfigured its drainage pattern, driving the dispersal and diversification of *Hypostomus*. The Miocene marine incursion was a strong barrier and its retraction triggered *Hypostomus* dispersal, increased speciation rate and ecological diversification. The timing of hydrogeological changes in La Plata Basin coincides well with *Hypostomus* cladogenetic events, indicating that the history of this basin has acted on the diversification of its biota.

## Introduction

During the Neogene (23 to 2.6 Mya), Neotropical river networks underwent a complex evolution due to marked changes in geomorphology and environmental conditions, most likely driving great biological diversification in freshwater fishes ^1–4^. However, considerable uncertainty remains about the precise links between the region’s geomorphological history and the evolutionary origin, geographical distribution and diversification pattern of modern Neotropical fish fauna. Among Neotropical freshwater fishes, armored catfishes of the family Loricariidae represent an excellent model to study the effects of landscape evolution on fish lineage diversification. This family is the most species-rich among Siluriformes and has members in all Neotropical basins. Over more than a quarter-century, investigations of the molecular phylogeny of loricariids have resulted in important discoveries and changes in their classification ^5–10^. However, there are still several unresolved phylogenetic relationships in this family, such as in lineages with fast diversification, or due to the under-representation of species coming from particular river basins. These remaining challenges limit the investigation of important questions regarding on the evolution of these Neotropical freshwater fishes.

The genus *Hypostomu*s is one of the most diverse and widespread members of the family Loricariidae, and inhabits most of the Neotropical Realm ^11^. The phylogeny and systematics of this group has attracted much interest ^10,12–17^. Particularly, in La Plata Basin (which includes the rivers Paraguay, Paraná, Uruguay and Río de La Plata) this genus is represented by more than 50 species, becoming one of the most species-rich genera of this large basin ^11,18,19^. This feature is very relevant since La Plata Basin is the third most speciose basin in South America, after the Amazon and the Orinoco basins ^20^. Thus, *Hypostomus* is an excellent model to comprehend the events that have given rise to the current diversity of fish species in La Plata Basin. However, understanding the causes of this species richness requires detailed species-level knowledge of geographic distribution, ecologically relevant traits and phylogenetic relationships.

Understanding why members of a clade have dispersed to some places and not to others is essential to recognize large-scale patterns of clade distribution. Historical biogeography studies have become narrowly focused on using phylogenies to discover the history of connections among regions. The geographical distribution of a clade will be determined by (i) the ancestral ecological niche and the opportunities for ecological niche shifts that are afforded to species by their geographical locations; (ii) the ancestral distribution area and the limitations (both physical barriers and/or biotic conditions) to dispersal; and (iii) the amount of time during which niche shifts and dispersal could have occurred ^21^.

For freshwater fishes, distribution patterns and dispersal dynamics must be interpreted in the context of the geological and geographic histories of river basins ^20^. Unlike vagile terrestrial organisms, the ability of strictly freshwater fish to move between drainages is limited by the hydrological connectivity. To understand how the complex geomorphological history of La Plata Basin may have affected the geographical distribution of fish species, it is crucial to infer the phylogenies of widely distributed fish lineages. These phylogenies contain information to trace past physical connections and disconnections within and between freshwater habitats^1,3,22^. Paleogeographically structured biogeographic patterns may inform about species divergence times especially when no fossils have been assigned to the clade^23^.

### Geomorphological changes in La Plata Basin

Approximately 30 Mya, the Michicola Arc uplifted generating the division between the proto Amazonas-Orinoco and La Plata Basin^24^. However, the boundary between both basins has been locally remodeled since then, providing dispersal opportunities between basins for freshwater organisms. For instance, ^25^ proposed a notable river capture event between the Pilcomayo River (La Plata Basin) and the Upper Rio Grande Basin (Amazon Basin) 3.5 Mya, which shifted the divide between the two basins farther north. Also, ^26^ suggested that the Amazon-Paraguay divide is a complex semipermeable geomorphological structure shaped by the Andean orogeny, the megafan environment, and climatic factors, which allowed the migration of several taxa from tributaries of the Amazon to La Plata Basin.

Marine transgressions occurred at least twice in La Plata Basin during the Miocene 15-5 Mya. The first maximum flooding event occurred between 15 and 13 Mya ^27,28^ and the second between 10 and 6.8 Mya ^29–31, 32^. The two marine transgressions covered most of the current Lower Paraná River Basin ^28^, with a considerable impact on the distribution of freshwater organisms.

The pattern of tributaries of La Plata Basin has undergone numerous modifications through time. Although there is no consensus on the different positions taken by the course of the Paraná River over time, ^33,34^ proposed that in its initial configuration, the Upper Paraná River was pouring its waters directly into the Uruguay River. Later, about 7 Mya, the lower section of the Upper Paraná changed its course towards the west and connected to the Paraguay river (forming the Lower Paraná), resulting in its current configuration. However, according to ^35^, the Upper Paraná River was initially turning eastward and headed towards the Patos lagoon (Brazil). Then it took its current configuration about 2.5 Mya. The current hydrographical pattern of the Paraná and Uruguay rivers indicate that they are exclusively connected via the Río de la Plata estuary. However, several authors ^36–38^ have proposed contemporaneous water pathways between the Middle Paraná and Middle Uruguay rivers during flood periods.

On the east side of the La Plata Basin, river headwater capture events between the Upper Paraná and the coastal rivers of southern Brazil were documented ^39^. According to ^40^, these capture events have been very numerous between 28-15 Mya and may explain the origin of the fish diversity of the coastal rivers of southern Brazil through dispersal processes ^41^.

In this work, we use the species-rich catfish genus *Hypostomus* (Loricariidae) as a model group to determine from where and when Amazonian species colonized La Plata Basin, and how they diversified in this basin that has undergone intense orogenic changes. We addressed these questions by inferring a time-calibrated phylogeny for *Hypostomus* with an almost complete representation of species inhabiting La Plata Basin. Using our chronogram together with dated changes in the hydrological pattern of the basin such as the marine incursions, we reconstructed the ancestral distribution ranges and estimated the ancestral preferred habitats for *Hypostomus* species, allowing us to propose an integrated view of the history of diversification of this genus in La Plata Basin.

## Materials and methods

### Taxon sampling and data collection

We analysed 52 species of *Hypostomus* (32 of which were collected in La Plata Basin), representing most of the species described to date in La Plata Basin and covering all the rivers of this basin. Specimens were euthanized immediately after collection in a solution containing a lethal dose of Eugenol (clove oil). Tissue samples for genetic studies were preserved in 96% ethanol and frozen at −20 °C. The voucher specimens were fixed in formalin 4% and deposited at MHNG, IPLA, and MACN (the institutional abbreviations are as listed in Ferraris, 2007) and at Fundación de Historia Natural Félix de Azara, Buenos Aires, Argentina (CFA-IC). We included five species of closely related genera as outgroups.

### Ethics statement

In argentina, fish were collected with the permission of the local authorities: Ministerio de Ecologíia y Recursos Naturales Renovables de Misiones (Disp. 2015 and 2016); Ministerio de Producción y Ambiente de Formosa (N ° 040/2015); Secretaría de Medio Ambiente y Desarrollo Sustentable de Santa Fe (Res. 081/2015); Ministerio de Asuntos Agrarios de Buenos Aires (Res. 355/10); Dirección de Fauna y Areas Naturales Protegidas de Chaco (Cons. Aut. 2011); Dirección de Flora, Fauna Silvestre y Suelos de Tucumán (Res. 223/2015); Secretaría de Medio Ambiente y Desarrollo Sustentable de Salta (Res. 091/2015) and Dirección General de Bosques y Fauna de Santiago del Estero (Ref. 17461/2015). Our Institutions do not provide the statements on ethical approval of the study. Notwithstanding, this study was approved by the National Council of Scientific and Technical Research of Argentina (exp. 7879/14) and it is a requirement of this institution to follow the guidelines of its “Comité de Etica” (https://www.conicet.gov.ar/wp-content/up-loads/OCR-RD-20050701-1047.pdf) and its biological sampling guide (https://proyectosinv.conicet.gov.ar/solicitud-scientific-collection/). Also, fish handling during sampling was performed following guidelines of the UFAW Handbook on the Care and Management of Laboratory Animals (http://www.ufaw.org.uk). Fish were anesthetized and killed using water containing a lethal dose of eugenol (clove oil). Other used specimens and tissues are deposited in Museu de Ciências e Tecnologia da PUCRS; Muséum d’histoire naturelle - Ville de Genève; and Laboratório de Ictiologia e Pesca of the Universidade Federal de Rondonia (Porto Velho, Brazil).

### DNA isolation, PCR, and sequencing

Genomic DNA was extracted using the salt-extraction protocol^42^. Polymerase chain reaction (PCR) was used to obtain two mitochondrial DNA (mtDNA) markers: 592 bp of Control Region (D-loop) and 826 bp of Cytochrome Oxidase Subunit I (COI), and partial sections of three nuclear markers ^43^ : 1345 bp of Recombinase Activating gene-1 (RAG1), 1059 bp of Hodz3 gene and 1144 bp of HAMzbtb10-3 gene. Hodz3 (*Hypostomus* ortholog of the Odz3 gene in vertebrates) encodes a transmembrane protein with a conserved role in eye development in vertebrates ^44^. The anonymous marker HAMzbtb10-3 is most probably a fragment of intron number 3 of the gene zbtb10 (zinc finger and BTB domain containing 10), as identified in the genome of the catfish Ictalurus punctatus (Source:NCBI gene;Acc:108257261). According to SINE Base (http://sines.eimb.ru/), it includes a sequence of 313 bp that is a putative SINE of the Ras1 type ^45^.

Each PCR amplification was performed in a total volume of 50 µl, containing 5 µl of 10x Taq DNA Polymerase reaction buffer, 1 µl of deoxyribonucleoside triphosphate (dNTP) mix at 10 mM each, 1 µl of each primer at 10 µM, 0.2 µl of Taq DNA Polymerase equivalent to 1 unit of Polymerase per tube, and 1 µl of genomic DNA. Cycles of amplification were programmed as follows: (1) 3 min. at 94 °C (initial denaturation), 30 sec. at 94 °C, (3) 30 sec. at 52–56 °C, (4) 1 min. at 72 °C, and (5) 5 min. at 72 °C (final elongation). Steps 2 to 4 were repeated 40 times. All used primers are listed in Table 2. The PCR products were purified and sequenced by the company MAGROGEN (Korea). A full list of analysed specimens and their respective sequence GenBank accession numbers is given in Table S1.

### Sequence alignment, congruence among markers and phylogenetic analysis

The editing and alignment of the sequences were performed using BioEdit v.7.0.1 ^46^. For each marker and for the alignment of five concatenated markers, we implemented JModelTest2 v.2.1.10 ^47^ to estimate appropriate substitution models with the Akaike information criterion (AIC).

To assess whether the five used loci could be combined in a joint analysis, we used the Congruence Among Distance Matrices (CADM) test ^48^ implemented in the R library ape ^49^. A Kendall’s coefficient of concordance W value of 0 signifies complete incongruence among loci, while a value of 1 indicates complete con^-^ cordance. The null hypothesis of complete incongruence among DNA sequence distance matrices was tested with 999 permutations. We also assessed congruence among gene phylogenies by running individual gene tree analyses and calculating gene (gCF) and site (sCF) concordance factors in IQtree ^50^ and following ^51^. In these analyses the gCF for each branch is the fraction of decisive gene trees concordant with this branch and the sCF for each branch is the fraction of decisive alignment sites supporting that branch.

We used the concatenated matrix, partitioned by gene, to perform two phylogenetic reconstruction methods. First, maximum likelihood (ML) reconstructions were performed using RAxML ^52^. Confidence values for internal edges of the ML tree were computed via bootstrapping ^53^, based on 1,000 replicates. Second, a Bayesian Inference (BI) analysis was conducted in MrBayes 3.2 ^54,55^. Four chains were run simultaneously (three heated, one cold) for 10 million generations, with tree space sampled every 100th generation. After a graphical analysis of the evolution of the likelihood scores, the first 300,000 generations were discarded as burn-in. The remaining trees were used to calculate the consensus tree. Both reconstruction (ML and BI) methods were run on the CIPRES Science Gateway computing cluster ^56^.

### Time-calibrated Tree

A time-calibrated phylogenetic reconstruction based on the concatenated data was performed using BEAST v.1.7.5 ^57,58^. We used a different substitution model for each marker as selected through Jmodeltest2, with the Yule speciation process and a log normal-relaxed molecular clock. The analy^-^ sis was run for 20 million generations and sampled every 1,000th generation. The convergence of the runs and ESS values were investigated using Tracer v.1.7.5 ^59^. A conservative burn-in of 25% was applied after checking the log-likelihood curves. Finally, a maximum credibility tree with median ages and the 95% highest posterior density (HPD) was subsequently generated using TreeAnnotator v.1.7.5^58^. Due to the absence of an appropriate fossil record, we used two calibration time points based on dated hydrogeological changes. The first point was proposed by ^3^ and consists in attributing 8 Mya to the speciation between *Hypostomus hondae* and its closest relative, *H. plecostomoides. Hypostomus hondae* is distributed only in the Lago Maracaibo and Magdalena basins, while *H. plecostomoides* is known only from the Orinoco basin and some localities in the upper Amazon. These distribution patterns match the vicariant episode that separated Lago Maracaibo system from the Amazon and Orinoco basins 8 Mya ^24^. We incorporated a second and new calibration point. Our phylogenetic analyses indicate that *H. formosae*, distributed in the Pilcomayo River (draining into La Plata Basin), is the closest relative to *Hypostomus* sp. BO14-052, which comes from the Rio Grande, a tributary of the Rio Madeira Basin (draining into the Amazon Basin). Since these distribution patterns match the headwater capture proposed by ^25^ 3.5 Mya (Table 1), it is reasonable to attribute this age to the allopatric speciation event that gave rise to *H. formosae* and *Hypostomus* sp. BO14-052.

**Table 1.** Documented past hydrogeological events in La Plata Basin.

| Hydrogeological events | Mya | References |
| --- | --- | --- |
| Boundary displacements between the proto Amazonas-Orinoco and La Plata Basin | 30-20 | 24 |
| River headwater capture events between the Upper Paraná and the coastal rivers of southern Brazil | 28-15 | 26 |
| First maximum flooding event of marine incursion in La Plata Basin | 15-13 | 28 |
| Second maximum flooding event of marine incursion in La Plata Basin | 10-6.8 | 28,29,31,32 |
| Boundary displacements between Paraná and Uruguay rivers | 7-2.5 | 33,35,37 |
| River headwater capture events between the Rio Grande and Pilcomayo River | 3.5 | 25 |

**Table 2.**
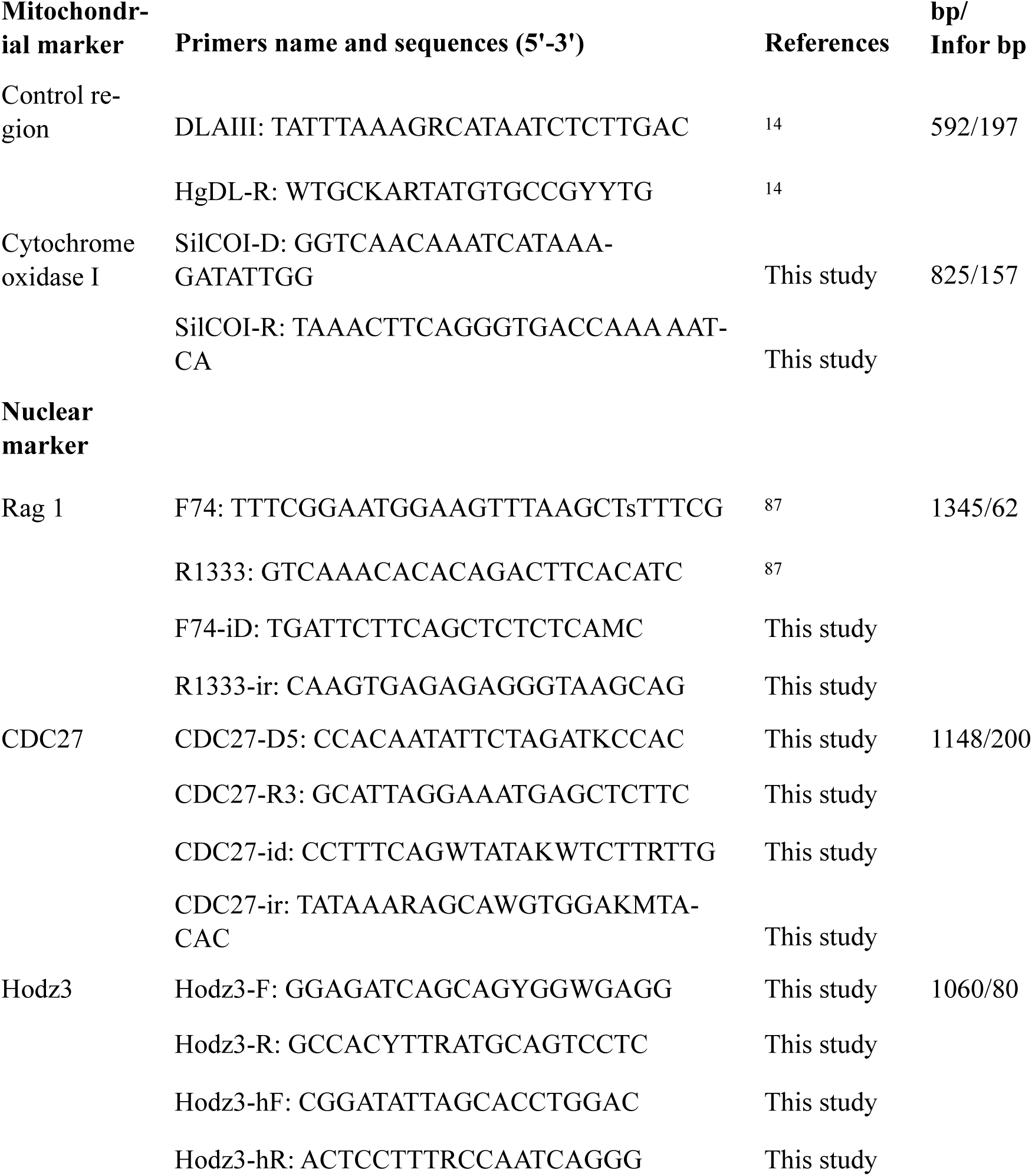
Primers used to amplify the five markers, the sequences length of the amplified DNA fragment (bp) and the number of phylogenetically informative bp (Infor bp).

### Inference of Ancestral Areas

To infer the biogeographic history of *Hypostomus* in La Plata Basin, we used the Dispersal-Extinction-Cladogenesis model (DEC)^60^ in the software RevBayes^61^.

As our interest focuses on La Plata Basin, we have decided to simplify the hydrological ecoregions of the Amazon, Orinoco, Guiana Shield, and Pacific Coastal drainages in only one region, called C (Fig.1). For La Plata Basin, we used the hydrological ecoregions proposed by Abell et al. (2008): Upper Paraná Basin (E), Paraguay Basin (B), Middle and Lower Paraná Basin and the Endorreic Andean basins (A), and Uruguay Basin (D). We also used other nearby regions such as the Northeastern Brazilian drainages, which also encompass the São Francisco River (G) and the Southeastern Brazilian drainages (F). Distribution areas of the *Hypostomus* species analysed here were taken from many primary-source publications and the collected specimens of CFA, MCP, MACN, MHNG.

In our analyses, we constrained rates of dispersal between ecoregions that are separated (non-adjacent) by one intercalated region to 1/10 of the dispersal rate between adjacent ecoregions. Moreover, due to Miocene marine transgression (10-6.8 Mya) that flooded the Middle and Lower Paraná section with saltwater, and because *Hypostomus* species live in freshwater, we excluded dispersal events crossing the Middle and Lower Paraná Basin (A) during the time window of this marine transgression [time-stratified constraint (MT)]. The tables of pairwise area connectivity constraints are given in Fig. S3 for cases without marine transgression (MT-) and with marine transgression (MT+). As a null model (non time-stratified model), we also ran the same analysis disregarding the changes in the connectivity among ecoregions during the marine transgression. Bayes factor was used for the comparison between both models ^23^. Both models were run based on default values as described in the RevBayes tutorial (http://revbayes.github.io/tutorials.html), with a total of 10,000 generations. Since the range size o the most widespread species in our phylogeny covers three ecoregions, we constrained the ancestral range reconstruction to include a maximum of three discrete ecoregions.

### Ancestral habitat reconstruction

Habitat preference for each species was classified according to one or more of the following five freshwater habitat categories, taken from ^62^. Habitat preference was taken from the original species description, catalogues, checklists and the authors’ personal field notes:

1. Upland streams and small rivers in Andean piedmont: swiftly flowing, well oxygenated and rich in dissolved minerals. The substrate comprises rocks and stones; usually between 800 m and 1200 m above sea level.
2. Upland streams and small rivers in shield escarpments: also fast flowing, well oxygenated, but carrying little suspended sediments and dissolved minerals; generally between 200 m and 1000 m above sea level.
3. Lowland *terra-firme* streams and medium rivers lying above the extent of seasonal flooding and over Tertiary formations; usually below 200 m contour. Water with suspended sediments and nutrients and tea-like blackwater coloration. Flow rates are typically low; the substrate is comprised of sand and clay.
4. Deep river channels: deep, swiftly flowing rivers with a seasonal flood pattern; typically exceeding around 3-5 m maximum depth during a typical low water. Turbid water, rich in suspended sediments and nutrients, which imparts a *café au lait* coloration.
5. Tidal estuarine systems with freshwater.

To infer ancestral habitat (discrete traits) across time in the *Hypostomus* tree, we ran the rayDISC function in the R package corrHMM ^63^ which is a maximum-likelihood method that allows multi-state characters and polymorphic characters. Species showing polymorphic habitat states were assigned a priori to occur in distinct habitats. We ran three different models: (i) one-parameter equal rates (ER) model, in which a single rate is estimated for all possible transitions; (ii) symmetric (SYM) model, in which forwards and reverse transitions between states are constrained to be equal; and (iii) all rates different matrix (ARD) model, in which all possible transitions between states receive distinct values. The corrected-Akaike information criterion (AICc) was assessed to choose the best model representing the evolution of habitat preference in *Hypostomus* phylogeny.

### Diversification through time

Analyses based on the time-calibrated phylogenetic tree were performed to quantify the evolutionary tempo and mode of diversification in *Hypostomus* within La Plata Basin. Firstly, using our time-calibrated tree we created lineages through-time (LTT) plot implemented in the R package ape^49^ based on only *Hypostomus* from La Plata Basin. To do this we pruned from the *Hypostomus* phylogenetic tree the specimens not included in the desired dataset. Second, to quantify historical rates of species accumulations within La Plata Basin (i.e., tests that diversification has changed over time), we compared our LTT plot with 1000 simulated trees under a constant-rate birth-death model, with identical root age and number of terminals. The simulations were performed using the function ‘ltt.plot’ (script developed by Hallinan & Matzke, available at http://ib.berkeley.edu/courses/ib200b/labs/lab12/rbdtree.n3.R), and values of speciation and extinction rates to feed this function were estimated using the R package ‘diversitree’ v. 0.9.10 ^64^. Louca and Pennell ^65^ revealed that an infinite number of pairs of speciation and extinction rates could give rise to any given LTT plot, and it is thus unclear how to determine the correct diversification rates. Therefore, phylogenies based only on extant species may not provide reliable information about past dynamics of diversification. Then, to circumvent the potential problem that underlies LTT plots, we followed Louca and Pennell’s ^65^ recommendation by calculating the ‘pulled diversification rate’ (PDR). PDR is a variable that is based on a set of congruent diversification scenarios that are compatible with a single LTT, and describes the relative change in diversification rate through time^65^. To do this analysis, we used the function ‘fit_hbd_pdr_on_grid’ of the R package ‘castor’ ^66^. Also, we graphically represented the rate of speciation of the *Hypostomus* genus only within La Plata Basin. This analysis shows the number of speciation for every 1 Mya time windows. We plotted each principal clade separately and the total cladogenetic events for the entire basin. Unlike most of the LTT plots used, our graph is not cumulative.

## Results

### Molecular data set

The final concatenated alignment comprised 4966 bp, including two mitochondrial (D-loop and COI) and three nuclear markers (RAG1, Hodz3, and HAMzbtb10-3). A total of 52 *Hypostomus* species were represented in the alignment, 32 of which are distributed in La Plata Basin. The concatenated data set has only 1.7% of missing character data. Kendall’s coefficient of concordance (W) rejected the incongruence among the matrices of the five loci, globally (W = 0.796, *p* = 0.003). Thus, the results of the CADM test indicated that all loci could be combined in a single phylogenetic analysis. The calculated gCf and sCF for each branch of the concatenated tree are shown in Fig. S1. Table S2 shows the best nucleotide substitution models for each partition evaluated in JModelTest2 and used in the phylogenetic analyses.

### Phylogenetic relationships

Phylogenetic analyses based on the five loci provided substantial resolution of relationships among species of *Hypostomus* (Fig. 1). Both maximum likelihood tree (ML) and Bayesian Inference (BI) retrieved the same topology, with well-supported clades [>70 bootstraping (BS) and >0.8 pos^-^ terior probability (PP)]. Our results supported the organization of *Hypostomus* into four groups, named D1, D2, D3 and D4 (all of them with BS=100 and PP=1), as initially proposed by Montoya-Burgos ^3^, retrieved by Cardoso ^14–16^ and recently by Jardim de Queiroz ^43^. However, our analyses showed a different relationship among these clades (D1(D2(D3+D4))) with high BS and PP values. Clade D1, forming the *H. cochliodon* group, comprised only one species inhabiting La Plata Basin (*H. cochliodon*). Clade D2, comprising the *H. plecostomus* group, contained 12 species from La Plata Basin. Clades D3 and D4, forming the *H. regani* group, were comprised of 19 species from La Plata Basin.

**Fig. 1.**
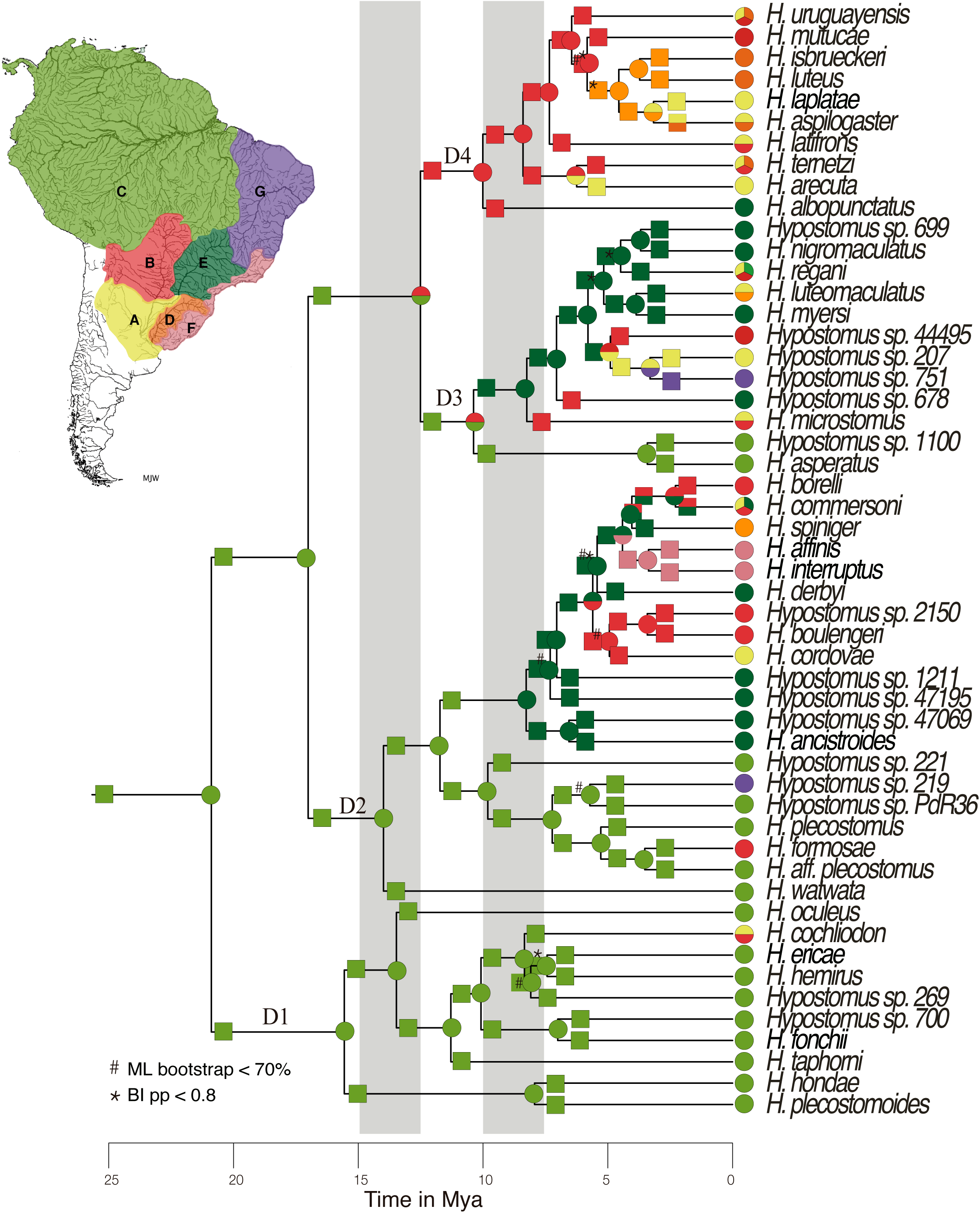
Time-calibrated multilocus phylogeny for the *Hypostomus* genus, illustrating ancestral areas estimations performed using RevBayes analysis. For the ecoregion codes and the D1 – D4 clades, see text. Grey vertical bars are the maximum flowing events of the Miocene Marine ingression. Lower bootstrap values and posterior probabilities are indicated.

### Relaxed clock, ancestral area and ancestral habitat estimation

According to our time calibrated tree reconstruction, *Hypostomus* started diversifying about 20.89 Mya (14.54-28.02 Mya 95% HPD). The ancestral range inference based on RevBayes shows that the genus originated somewhere in the Amazon-Orinoco-Guyane (hydrological ecoregion C in Fig. 1), and occupied upland streams and small rivers in shield escarpments (habitat 2 in Fig. 2; see also Table 3 that shows the AICc values to choose the best model used in the habitat inference).

**Table 3.** Nucleotide substitution models for each partition as evaluated by JModelTest and models used in the Maximum Likelihood (ML) and Bayesian Inference (IB) analyses.

| **Marker** | **JModelTest best model** | **Model used in Beast** | **Model used in MrBayes and RAxML** |
| --- | --- | --- | --- |
| Control Region | TIM2+I+G | HKY+I+G | GTR+I+G |
| Cytochrome oxidase I | TVM+G | GTR+G | GTR+I+G |
| Rag 1 | TVM+G | GTR+G | GTR+I+G |
| HAMzbtb10-3 | TVM+G | GTR+G | GTR+I+G |
| Hodz3 | GTR+I+G | GTR+I+G | GTR+I+G |

**Fig. 2.**
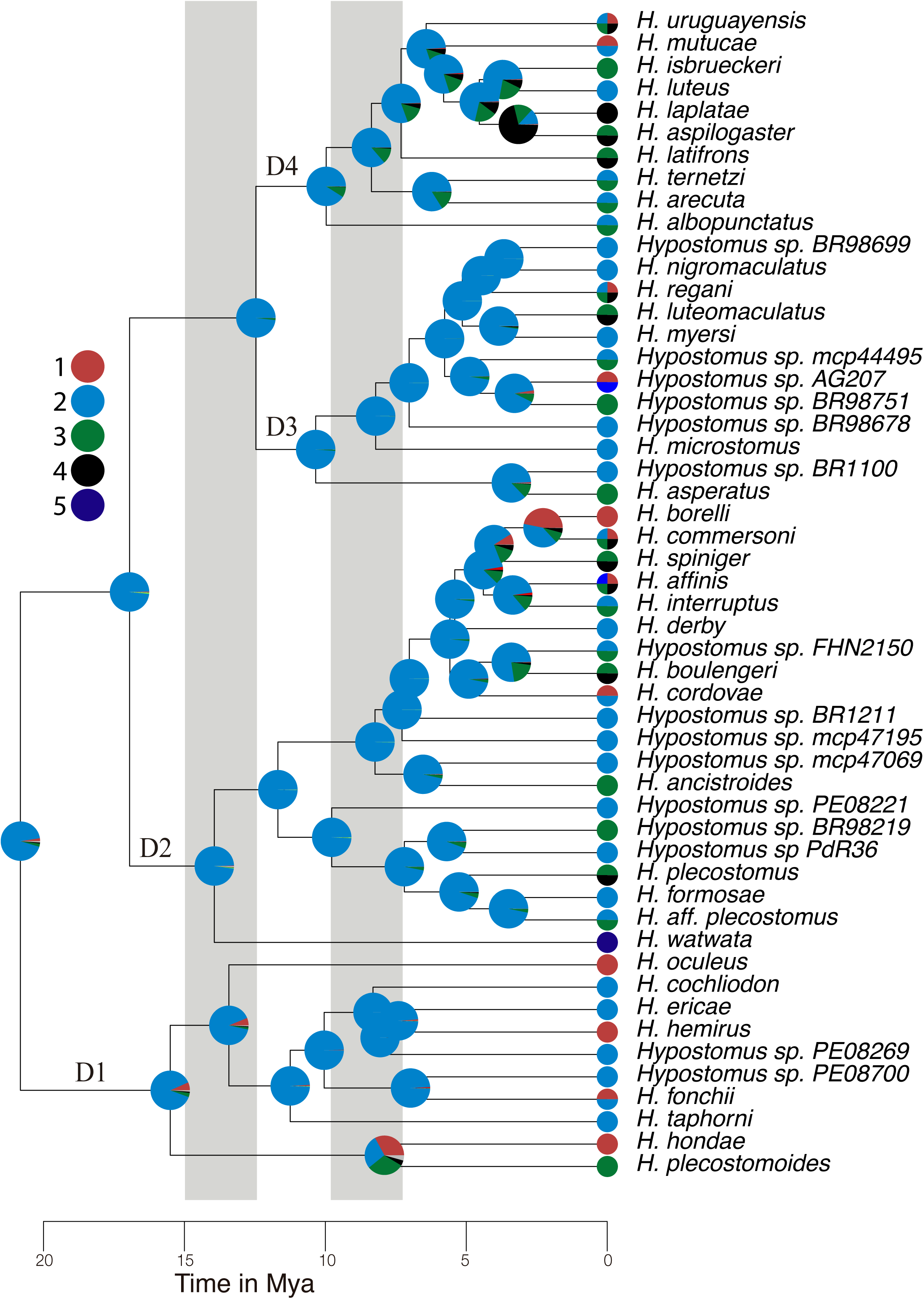
Time-calibrated multilocus phylogeny for the *Hypostomus* genus, illustrating ancestral habitat inferences performed using a Bayesian discrete phylogeographic approach. Numbers next to branches show their posterior probability. Habitat preference codes: upland streams and small rivers in Andean piedmont (1), upland streams and small rivers in shield escarpments (2), lowland *terra-firme* streams and medium rivers (3), deep river channels (4), and tidal estuarine systems (5). Grey vertical bars are the maximum flowing events of the Miocene Marine ingression.

The earliest split event in the phylogeny of *Hypostomus* divided the group *H. cochliodon* (Clade D1) from the rest of the genus. The diversifications of the Clade D1 started 15.56 Mya (10.85-20.6 Mya 95% HPD) and occurred mostly in the Amazon-Orinoco-Guyane (ecoregion C in Fig. 1). In Clade D1, RevBayes analysis estimated a single dispersal event at 8.34 Mya (6.33-12.56 Mya 95% HPD) from ecoregion C to B and A along the branch leading to *H. cochliodon*. Most *Hypostomus* species in Clade D1 occupied upland streams and small rivers in escarpments (habitat 2 in Fig. 2).

The ancestor of Clade D2 was also hypothesized to have originated in the Amazon-Orinoco-Guyane, approximately 17.01 Mya (11.86-22.78 Mya 95% HPD). The first split in clade D2 generated on one side *Hypostomus watwata* (from coastal Guyana region) and a large clade including 12 species inhabiting the La Plata Basin. This later clade shows on one side *H. formosae* is nested within a lineage of Amazon-Orinoco-Guyane species; on another side 11 species are grouped in a lineage composed principally of species from La Plata Basin. The origin of the later lineage was estimated to 11.73 Mya (7.86-15.67 Mya 95% HPD) in the Upper Paraná (ecoregion E in Fig. 1) after a dispersal event from Amazon-Orinoco-Guyane ecoregion. Our analysis of ancestral habitat inference showed that this ancestor inhabited small rivers (habitat 2 in Fig. 2). Habitat shifts and dispersal from Upper Paraná to other rivers of La Plata Basin as well as to the southern coastal rivers of Brazil were estimated to be younger than 7.05 Mya.

Clade 3 originated in the Paraguay ecoregion (B) after a dispersal event from the Amazon basin, while Clade 4 appeared in the initial ecoregion C (Fig.1). Members of both clades where living in small rivers (habitat 2 in Fig. 2). The first diversification event within Clade 3 was inferred as a vicariance event (but may correspond to the closure of a dispersal portal) between Amazon-Orinoco-Guyane and Upper Paraná approximately 10.38 Mya (6.87-14.2 Mya 95% HPD). Clade 3 includes nine species from La Plata Basin. Most ancestors of these species had the Upper Paraná (ecoregion E in Fig. 1) as ancestral area and occupied small and medium-sized rivers (habitat 2 in Fig. 2). Within this clade, dispersal events to other regions of La Plata Basin started 8.25 Mya. A lineage from São Francisco River (ecoregion G) is nested within the Clade D3 with a splitting age of 3.31 Mya (1.66-5.22 Mya 95% HPD)

Clade D4 contains ten species from La Plata Basin. Its first diversification event started in the Paraguay River (ecoregion B in Fig. 1), in small and medium rivers, approximately 10.01 Mya (6.69- 13.72 Mya 95% HPD). Dispersal events to other rivers of La Plata Basin and shifts in habitat preference were estimated to have occurred very early in the history of this lineage, concomitant with the beginning of the diversification of the D4 lineage.

### Diversification through time

In this analysis we plotted the numbers of speciation events through time. The LTT plot of *Hypostomus* from La Plata Basin showed an exponential growth in the lineage accumulation over time, but with a significant upshift in the cumulative number of lineages after 5 Ma (Fig. 3). The simulated LTT plots rejected a model of constant net speciation (γ= 1.226787e-01) and extinction (µ=1.985507e-06). Until 7 Mya, the shape of the LTT plot is just under the 75% confidence interval of the simulated LTT data under the constant speciation. Then, its slope increases and become passes the upper 95% confidence level at −5 Mya, higher than expected (above the 99 % confidence intervals at −4 Mya). The pulled diversification rates (PDR) of birth-death models that best explains the phylogenetic tree of *Hypostomus* from La Plata confirmed the pattern observed with the LTT plot, indicating a relative increase of lineage diversification at 5 Mya (Fig 3). We plotted the numbers of speciation events through time only within La Plata Basin (Fig. S4). This figure indicates the number of splitting events for each clade in a time window of one Mya, as well as for the total speciation of the *Hypostomus* genus within La Plata Basin. We also plotted the trend line of each data. The speciation process within this basin peaked between 3 and 5 Mya, with a total of 12 speciation events in this time window. The trend line of the total data showed a positive tendence, thus the most speciation events were young. Our results also show the relative contribution of the three clades with species from La Plata Basin (D2, D3 and D4) in the species diversification in this basin (Fig. S4). The clades D2 and D3 showed also a trend line with positive tendence, strongly influencing the total data. Conversely, the clade D4 showed a trend line with almost no slope through time, thus the number of speciation events was constant within this clade.

**Fig. 3.**
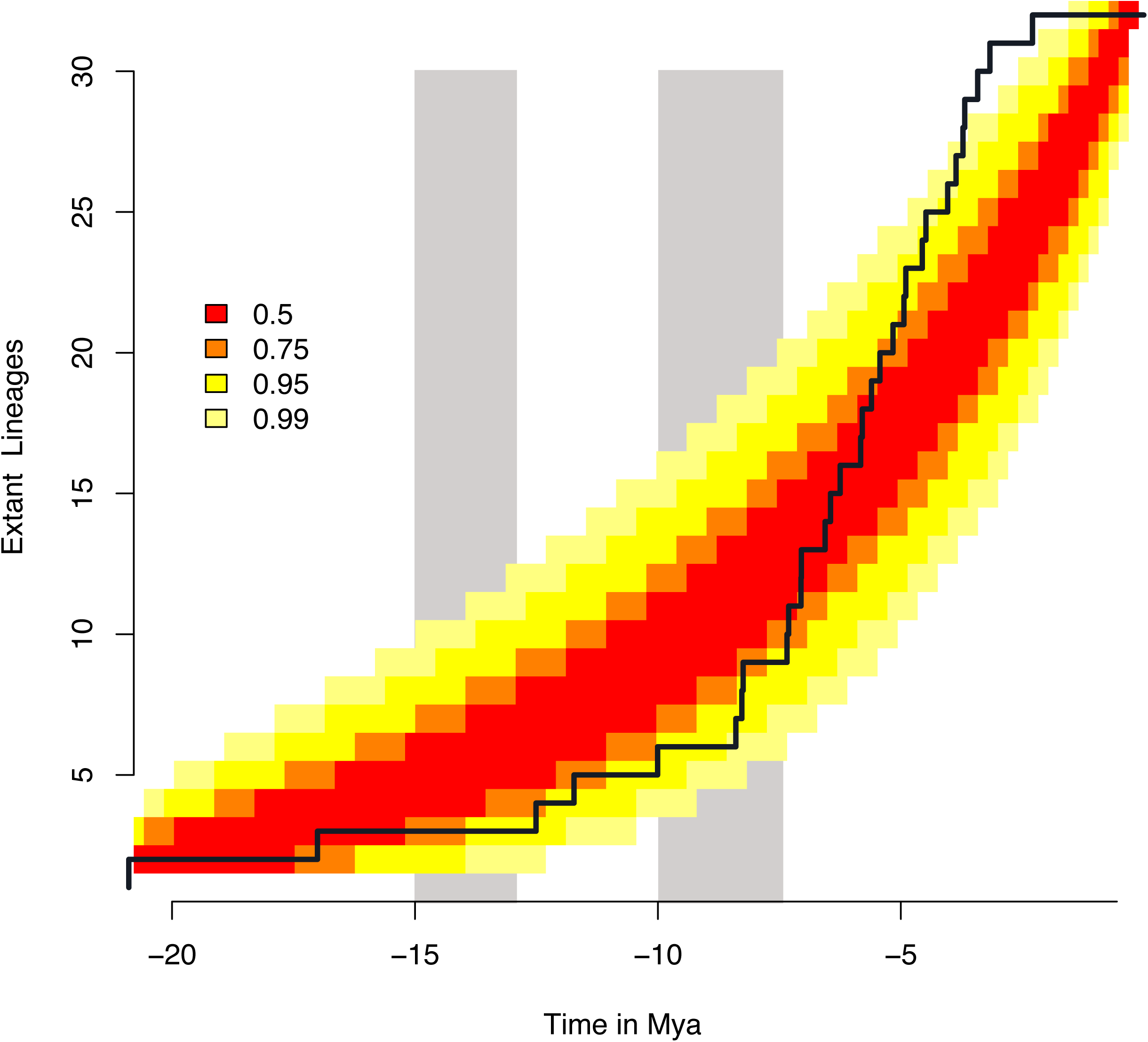
Tempo and mode of evolutionary diversification in *Hypostomus* within La Plata Basin. The black line shows the lineage through time (LTT) curve of species accumulation over time. Colors are the confidence intervals of a neutral model. Blue line is the Pulled Diversification Rate (PDR) over time. Grey vertical bars are the maximum flowing events of the Miocene Marine ingression.

## Discussion

In this study, we generated the most thoroughly-sampled, species-level phylogenetic hypotheses for the *Hypostomus* species inhabiting La Plata Basin, to date. The recovery of identical topologies under BI and ML methods based on five loci, with nodes supported with high bootstrap and PP values, suggests that strong phylogenetic signal is present in the data and that the relationships have been accurately reconstructed. Our robust *Hypostomus* phylogeny provides the opportunity to investigate the processes of evolutionary diversification in one of the most species-rich Neotropical fish genera. Previous studies included fewer taxa or were based on fewer molecular markers, resulting in trees with lower statistical support and/or with polytomies within the inter-species relationships ^3,8,10,14^. Nevertheless, the main lineages within the genus are similar to those found in ^3,10,14–16,43^, where four main Clades D1, D2, D3, and D4 were identified.

The South American fish fauna is the most diverse freshwater fish assemblage in the world and currently comprises more than 5000 described species, with many more still undescribed ^67^. In their phylogeny, many South American fish groups show sister clade relationships between current basins, most likely reflecting major hydrogeological changes in the continent ^68^. Indeed, there is substantial evidence that South America has undergone marked changes over the last 90 Mya, including marine incursions, weathering of ancient shields, uplift of the Andes and of palaeo-arches, significant reconfigurations in the size and drainage patterns of the principal river basins ^24,69^

Our phylogenetic results show that the *Hypostomus* species inhabiting La Plata Basin today do not form a monophyletic group. This result indicates that the species richness in this basin has not been entirely formed through the *in situ* diversification of a single ancestor, but is attributed in part to multiple colonization events. ^70^ analysed several freshwater fish clades under BPA and showed that the fish species pool of modern La Plata Basin were assembled during the rise of the Michicola Arch ∼24 Mya. Our chronology places the origin and the first split within the *Hypostomus* genus in the early Miocene (∼20 Mya), and geographically outside from the current La Plata Basin. The polyphyletic origin of *Hypostomus* species from La Plata Basin, combined with the middle Miocene age (∼17 Mya) of the three independent colonization events, imply that *Hypostomus* is not part of the ‘Old Species of La Plata Basin’. However, this younger age of first colonisation agrees with others studies on *Hypostomus* ^3,10,14^ and also on other subfamilies of Siluriformes (i.e, Callichthyinae ^71^).

Interestingly, our results suggest that the ancestors of the *Hypostomus* lineages from La Plata Basin colonized this basin via the Upper Paraná and Paraguay rivers and their descendants remained confined in this region until the end of the marine incursion which operated as a barrier in the Lower Paraná River (∼ 7 Mya).

The predictions of the principle of vicariance assume that multiple lineages will be affected by the raising of the putative barrier, giving origin to several sister-species pairs with similar ages ^72^. Conversely, the disappearance of a barrier does not imply the opportunity to disperse for all taxa. Certain geomorphological and climatic processes operating on continental platforms displace barriers to organismal dispersal and probably affect all three processes of macroevolutionary diversification: speciation (species lineage splitting), dispersal (species geographic range expansion), and extinction (species termination) ^73^. However, the dispersal also depends on the species ancestral ecological niche and its opportunities for ecological niche shifts to live in new geographical locations.

The marine incursion was an obvious dispersal barrier during late Miocene; however, its impact on freshwater fish dispersal may have persisted, at least partially, after the marine water receded, because such events can permanently alter soil properties and river water characteristics ^74^. Therefore, the dispersal of the *Hypostomus* lineages, after this marine incursion, depended on the ancestral ecological niche. The ancestral habitat and the geographical starting point for dispersal of all *Hypostomus* ancestors of La Plata Basin were estimated by our analyses of area and ancestral habitat inferences as follows. The ancestors of Clades D2, D3 and D4 occupied upland streams and small rivers in shield escarpments (habitat 2). The adaptation to lowland *terra firme* streams and deep river channels (habitats 3 and 4) and the colonization of other ecoregions started ∼6.5 Mya, short after the regression of the sea. This result indicates that the ancestral species that crossed the lower parts of La Plata Basin after marine water receded had the correct habitat preference for dispersal through this basin, and thus were adapted to live in larger rivers like Lower Paraná. In contrast, the environmental effect of the marine incursion and the ancestral habitat preference of the Clade D1 could explain why the *H. cochliodon* group showed a very low rate of diversification and dispersion in La Plata Basin. Clade D1 gave rise to a single species that occupies highland streams and small rivers in escarpments in this basin, which it colonized later (∼8 Mya) than the other clades. In fact, the *H. cochliodon* group (formerly the genus *Cochliodon* Kner) and the genus *Panaque* Eigenmann are capable of eating wood, an exceptional diet among fishes ^75^ and have large, spoon-shaped teeth. In contrast, the other *Hypostomus* species inhabiting in La Plata Basin have the typical diet of loricariids, they are algivorous or detritivorous. The habitat preference and the diet differentiation reflects a particular ecological niche of La Plata Basin representatives of Clade D1 that could have played a key role in delaying the dispersion of this group in La Plata Basin.

Once the Upper Paraná was occupied by *Hypostomus* ancestral species of Clades D2 and D3, their descendants could have disperse throughout the La Plata Basin after the period of sea level regression, a process opening speciation opportunities. Our analysis of LLT plots rejected a model of constant net speciation and extinction and showed an upshift of speciation rate after 5 Mya and other lines of evidence ^10^ suggest that the Paraná River acted as “biogeographical corridor” for the genus *Hypostomus* to colonize and speciate in other rivers belonging to La Plata Basin today (Paraguay, Uruguay and Río de la Plata). Even the LTT plots were strongly criticized recently ^65,76^, our LTT plot agrees with the PDR and the increase of the number of speciation events in our tree after 7 Mya. Consequently, the *Hypostomus* species of these rivers, which have a tropical origin, contrast with the xerophilous forests and temperate steppes (grassland and savannas) through which most of the rivers run. The effect of the Paraná River as biogeographical corridor of tropical freshwater-related biota towards temperate latitudes is a pattern described by numerous authors in relation to a number of diverse plant and animal groups ^77–79^. The existence of a relatively mesic microclimate on the riverbanks and the development of riparian humid forests and wetlands facilitate the dispersion of tropical floristic and faunistic species in temperate latitudes of the Plata Basin, in this way promoting new diversification.

La Plata Basin is a well-connected system today. Thus, in the absence of geographical barriers to dispersal, this basin may have promoted the mixing of populations at the regional level and reduced the likelihood of allopatric speciation. However, ancestral species of *Hypostomus*, subsequent to the multiple independent colonization events from the Amazon, diversified in La Plata Basin, showing both ecological and morphological diversification. The rate of species diversification within La Plata Basin markedly increased between 5 to 3 Mya. The current species diversity of *Hypostomus* in La Plata Basin is lower, but species are more phenotypically diverse than their congeners inhabiting other basins in the Neotropical Realm (as shown by ^10^. The heterogeneity of the water environment may be promoting the diversification observed in La Plata Basin. However, this basin was affected by numerous tectonic and climatic events during the last 5 Mya that did not leave obvious geographic features in the contemporary landscape. As shown for other regions of the Neotropics, our study suggests that the current drainage structure in La Plata Basin might be a poor predictor of *Hypostomus* biodiversity.

The colonization of La Plata Basin has not been carried out exclusively through the biogeographic corridor of Upper Paraná River. In the D1 clade, the *H. cochliodon* species from La Plata Basin is nested within the Amazon Basin species (Fig. 2), and our calibrated tree indicates a dispersion event from the Amazon to the Paraguay River (CB ecoregion) ∼ 8 Mya, by highland streams and small rivers (habitats 2). Moreover, the ancestral of clade D4, occupied the Paraguay River ∼ 10 Mya and the recent formation of *H. formosae* in the Paraguay River indicate anone other connection about 3.5 Mya ^26^ showed that the Paraguay and Upper Madeira rivers share most of same species. These authors suggested that the headwaters of some tributaries of the Amazon Basin and the Paraguay River (tributary of La Plata Basin) could interconnect during the rainy season, functioning as portals for the dispersal of fishes possessing the appropriate headwater ecology. However, this suggestion is less supported nowadays, as many of the cases mentioned by Carvalho & Albert ^26^ and others ^80^ have recently been recognized as distinct species by their genetics and/or morphology. For example, *Hoplias malabaricus*, considered widespread in South America, has been divided into 16 vicariant species ^81^. Inter-basin dispersal of fishes via exceptional portals followed by allopatric speciation has been documented in other regions of South America ^82,83^, but these flood events would have been older than thought. Because Neotropical ichthyology remains a dynamic field, with many new species described every year ^84^, the hypothesis of contemporary inter-basin dispersal portals opening during the rainy season will probably be challenged more time.

Our study of species diversification in the genus *Hypostomus* highlights the key role played by river captures. Here, we used a well-documented river capture between the Amazon and Paraguay River ^25^ to add a calibration point to the *Hypostomus* tree. Moreover, in our analysis of ancestral distribution range, we observed two signatures of river capture involving La Plata Basin. First, in Clade D2, we observed a vicariance event at ∼5 Mya between La Plata Basin and the coastal rivers of southern Brazil (ecoregion G). This split was very likely due to the past capture of the Upper Paraná headwaters by coastal drainages ^85^. Our results and previous ones ^3,10^ support the invasion by ancestral lineages of *Hypostomus* from La Plata Basin into the coastal rivers of southern Brazil. Furthermore, different lineages of Neotropical fish show similar patterns of dispersion and colonization. For instance, the time-trees of *Hoplosternum* and *Callichthys* show that both Callichthyidae catfish genera reached the coastal rivers of southern Brazil from the Upper Paraná, and that dispersal occurred from the Amazon to the Upper Paraná but apparently never vice-versa ^71^. In contrast, other Loricariidae (Neoplecostominae) and other freshwater fishes (Cynolebiini) had an ancestral origin in the coastal rivers of southern Brazil and dispersed later to La Plata Basin ^5,10,41,86^. According to ^40^, all these river capture events were very numerous between 28 and 15 Mya; however, our results and others ^5,10,41,71^ give evidence that more recent river captures have affected fish dispersal and diversity. Our results indicate a second river capture in Clade D3, in which a very well supported lineage shows a São Francisco species nested within La Plata Basin lineages. This result suggests that the Upper Paraná River was not only a “corridor” permitting the entrance of Amazonian species into La Plata Basin, but acted also as a passageway from the La Plata Basin to the São Francisco and to the coastal rivers of southern Brazil, at least for the *Hypostomus* genus.

Phylogenetic and biogeographical knowledge continues to be essential for the elucidation of species evolution, diversification, and the understanding of distribution patterns. However, it is necessary to note that the reconstruction of area cladograms do not depend solely on the history of geological connections among areas, but also on the history of connections among suitable habitats. The inference of a robust phylogenetic hypothesis further enhances the value of a given group of organisms as a model system for understanding Neotropical biodiversity. Our approach, incorporating different data sets (molecular, ecological, and geographical) to analyse the diversification history of a densely sampled fish lineage in the La Plata Basin (Fig. 4), has yielded thorough results for understanding the evolution, ancestral biogeographical ranges, and habitat preferences of *Hypostomus* species in this basin.

**Fig. 4.**
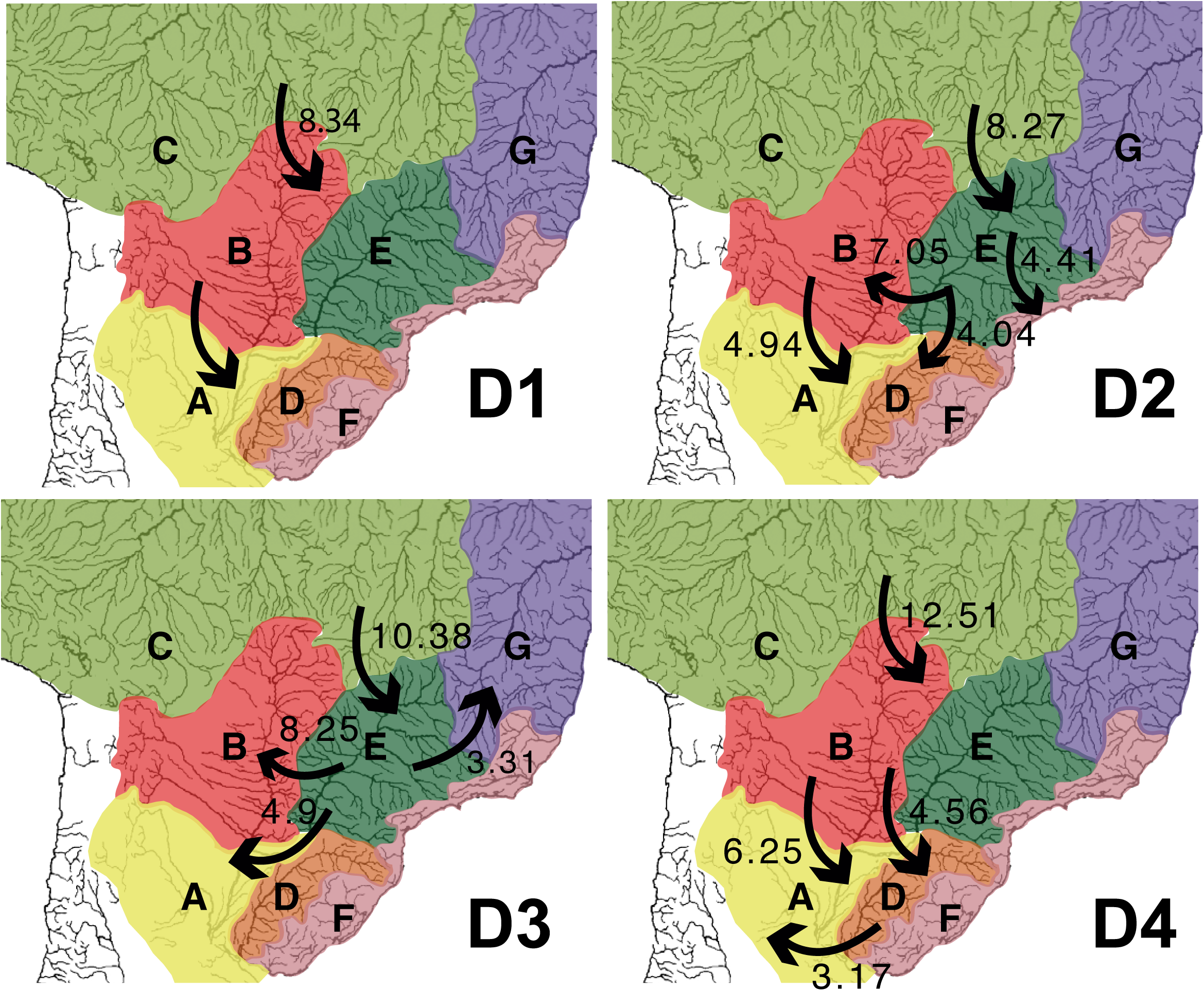
Graphic summary of the main colonizations in the evolutionary history of the genus *Hypostomus* in La Plata Basin. Major dispersals are shown with arrows. All dates are included and taken from our time-calibrated tree (Fig. 1).

## Acknowledgements

Numerous colleagues and students participated in the collection of materials and helped in this project, including Sergio Bogan and Juan M. Meluso (Fundación Félix de Azara); Raphael Covain and Sonia Fischer-Muller (Muséum d’histoire naturelle - Ville de Genève); Florencia Brancolini and Gustavo Somoza (Univer^-^ sidad Nacional de San Martín); Carlos Lucena (Museu de Ciências e Tecnologia da PUCRS); Lucila Protogino and Ariel Paracampo (Instituto de Limnología “Dr. Raúl A. Ringuelet”); Carlos Rivera-Rivera (University of Geneva); Adriana Almirón and Jorge Casciotta (Museo de La Plata) and Facundo Vargas (Fauna y Áreas Naturales Protegidas, Provincia del Chaco). We thank Nureen Ghuznavi (George Washington University) for English revision. Funding for this project was supported by PICT2014-0580 and PICT2017-1330 (YPC); Seed Money Latin America 2015 (JMB/YPC); BOLD-CONICET (YPC); Brazilian–Swiss Joint Research Programme 2015 (MJB/LJQ) and CNPq/Science without Borders (LJQ).

## Authors’ contributions

YPC and JIMB conceived the project; YPC, LJD, IAB, JIMB performed the laboratory work, conducted the fieldwork and collected the data with additional material from collaborators; YPC and LJD analysed the data; YPC drafted the manuscript; YPC, LJD, IAB, PEP and JIMB prepared the final version of the manuscript.

## Biosketch

YPC is broadly interested in the biogeography of Neotropical Region. This work represents a component of her PhD work at University of La Plata on the systematics and diversity of Loricariidae from La Plata Basin. LJD works on the evolutionary history of Neotropical fishes and on the processes underlying their tremendous diversity. This work is part of his PhD work at University of Geneva, Switzerland.

