## supplemental information for "Multilocus phylogeny and historical biogeography of *Hypostomus* shed light on the processes of fish diversification in La Plata Basin"

**SHORT RUNNING TITLE**: *Hypostomus* diversification in La Plata Basin

Yamila P. Cardoso[1]*†| Luiz Jardim de Queiroz[2]†| Ilham A. Bahechar[2]| Paula E. Posadas[1]| Juan I. Montoya-Burgos[2]

[1] Laboratorio de Sistemática y Biología Evolutiva, Facultad de Ciencias Naturales y Museo, Universidad Nacional de La Plata, Paseo del Bosque S/N, B1900FWA, La Plata, Buenos Aires, Consejo Nacional de Investigaciones Científicas y Técnicas, Argentina.

[2] Department of Genetics and Evolution, University of Geneva, 30 quai Ernest Ansermet, 1211, Geneva 4, Switzerland.

† These authors contributed equally to this work

Table S1. Full list of analysed specimens, collection code, GenBank accession numbers, collected locality, and habitat preferences code (see text).

| Species | Collection code | Genbank number (Dloop/COI/ Rag1/HAMzbtb10-3/Hodz3) | Location (Country, river system) | Habitat preferences |
|---|---|---|---|---|
| Hypostomus affinis | BR1254 | AJ318358/MK959853/ MK959930/MK959980/ MK960036 | Brazil, Agua Santa River | 1234 |
| Hypostomus albopunctatus | MCP37990 | AJ318379/MK959865/ MK959942/MK959992/ MK960048 | Brazil, Iguacu River | 23 |
| Hypostomus ancistroides | BR98696 | AJ318369/MK959857/ MK959934/MK959984/ MK960040 | Brazil, Rio Grande River Basin | 3 |
| Hypostomus arecuta | AG198 | JF290445/MK959840/ MK959916/MK959967/ MK960024 | Argentina, Middle Parana River | 3 |
| Hypostomus asperatus | BR974 | AJ318370/MK959854/ MK959931/MK959981/ MK960037 | Brazil, Bateia River | 3 |
| Hypostomus aspilogaster | DF023 | AJ318375/MK959860/ MK959937/MK959987/ MK960043 | Brazil, Rio Grande do Sul | 34 |
| Hypostomus borelli | BO14260 | MK959903/MK959848/ MK959925/MK959975/- | Bolivia, Pilcomayo River | 1 |

| | | | | |
|---|---|---|---|---|
| Hypostomus boulengeri | AR11609 | JX290097/MK959846/MK959923/MK959973/MK960030 | Argentina, Paraguay River | 34 |
| Hypostomus cochliodon | 1154 | JF290476/MK959839/MK959915/MK959966/MK960022 | Argentina, Middle Parana River | 23 |
| Hypostomus commersoni | YC09118 | MK959895/MK959883/MK959959/MK960012/MK960067 | Argentina, Rio de la Plata | 1234 |
| Hypostomus cordovae | AR111215 | KX852408/MK959844/MK959921/MK959971/MK960028 | Argentina, Segundo River | 12 |
| Hypostomus derby | YC10316 | JF290447/MK959885/MK959961/MK960014/MK960069 | Argentina, Middle Iguazu River | 1 |
| Hypostomus ericae | BR1013 | AJ318347/MK959849/MK959926/MK959976/MK960032 | Brazil, Maranhao River Basin | 23 |
| Hypostomus fonchii | PE08034 | MK959900/MK959868/MK959945/MK959995/MK960051 | Peru, Huecamayo River | 12 |
| Hypostomus formosae | AR11207 | JX290093/MK959845/MK959922/MK959972/MK960029 | Argentina, Paraguay River | 2 |
| Hypostomus hemirus | GY04333 | MK959898/MK959864/MK959941/MK959991/MK960047 | Guyane, Siparuni donwstream | 1 |
| Hypostomus hondae | VZ94 | AJ318348/MK959888/MK959956/MK960009/MK960064 | Venezuela, Muyapa River | 2 |
| Hypostomus interruptus | BR1161 | AJ412846/MK959851/MK959928/MK959978/MK960034 | Brazil, Ribeira do Iguape River | 23 |
| Hypostomus isbrueckeri | MCP41476 | AJ318376/MK959866/MK959943/MK959993/MK960049 | Brazil, Jacui River | 3 |
| Hypostomus laplatae | YC09031 | KX852412/MK959881/MK959957/MK960010/MK960065 | Argentina, Rio de la Plata | 4 |
| Hypostomus latifrons | PY8008 | AJ318378/MK959873/MK959950/MK960000/MK960056 | Paraguay, Salado River | 23 |
| Hypostomus luteomaculatus | AG200 | JF290467/MK959841/MK959917/MK959968/MK960025 | Argentina, Middle Parana River | 34 |

| | | | | |
|---|---|---|---|---|
| Hypostomus luteus | UR007 | AJ318374/MK959876/MK959953/MK960003/MK960059 | Brazil, Rio Uruguay River | 23 |
| Hypostomus microstomus | 1153 | JF290461/MK959838/MK959914/MK959965/MK960021 | Argentina, Middle Parana River | 2 |
| Hypostomus mutucae | Aqua9 | MK959894/MK959891/MK959920/MK960019/MK960072 | Brazil, Upper Paraguay River | 1 |
| Hypostomus myersi | YC10256 | AJ318355/MK959884/MK959960/MK960013/MK960068 | Argentina, Middle Iguazu River | 1 |
| Hypostomus nigromaculatus | Tib12 | AJ318355/MK959875/MK959952/MK960002/MK960058 | Brazil, Tibaji River | 2 |
| Hypostomus oculeus | PE08441 | MK959897/MK959871/MK959948/MK959998/MK960054 | Peru, Loreto | 1 |
| Hypostomus plecostomoides | VZ58 | AJ318349/MK959889/MK959955/MK960008/MK960063 | Venezuela, Apure/Masparo Bassin | 2 |
| Hypostomus plecostomus | SUJM047 | MK959905/MK959874/MK959951/MK960001/MK960057 | Suriname | 34 |
| Hypostomus regani | AG354 | MK959907/MK959843/MK959919/MK959970/MK960027 | Argentina, Middle Parana River | 1234 |
| Hypostomus sp. | AG207 | MK959908/MK959842/MK959918/MK959969/MK960026 | Argentina, Middle Parana River | 2 |
| Hypostomus sp. | BO14052 | MK959904/MK959861/MK959938/MK959988/MK960044 | Bolivia, Rio Grande | 2 |
| Hypostomus sp. | BR1100 | -/MK959892/MK960017/-/MK960073 | Brazil, Tocantins River | 3 |
| Hypostomus sp. | BR1211 | MK959906/MK959847/MK959924/MK959974/MK960031 | Brazil, Tiete River | 2 |
| Hypostomus sp. | BR98219 | AJ412834/MK959850/MK959927/MK959977/MK960033 | Brazil, Itapicuru River | 3 |
| Hypostomus sp. | BR98678 | AJ315764/MK959867/MK959944/MK959994/MK960050 | Brazil, Tiete River | 12 |

| | | | | |
|---|---|---|---|---|
| Hypostomus sp. | BR98699 | MK959896/MK959869/MK959946/MK959996/MK960052 | Brazil, Rio Grande | 2 |
| Hypostomus sp. | BR98751 | MK959901/MK959870/MK959947/MK959997/MK960053 | Brazil, Sao Francisco River | 2 |
| Hypostomus sp. | FHN2150 | MK959899/MK959872/MK959949/MK959999/MK960055 | Argentina, Bermejo River | 12 |
| Hypostomus sp. | MCP44495 | AJ412848/MK959855/MK959932/MK959982/MK960038 | Brazil, Upper Paraguay River | 2 |
| Hypostomus sp. | MCP47069 | AJ412839/MK959859/MK959936/MK959986/MK960042 | Brazil, Upper Parana River | 23 |
| Hypostomus sp. | MCP47195 | AJ412844/MK959852/MK959929/MK959979/MK960035 | Brazil, Upper Parana River | 2 |
| Hypostomus sp. | PdR36 | AJ412836/MK959856/MK959933/MK959983/MK960039 | Peru, Ucayali River | 2 |
| Hypostomus sp. | PE08221 | AJ412835/MK959858/MK959935/MK959985/MK960041 | Peru, Monzon River | 2 |
| Hypostomus sp. | PE08269 | -/MK959890/-/MK960018/- | Peru | 2 |
| Hypostomus sp. | PE08700 | -/MK959893/-/MK960020/- | Peru, Neshua River near Neshua | 2 |
| Hypostomus spiniger | YC09091 | MG457224/MK959882/MK959958/MK960011/MK960066 | Argentina, Uruguay River | 34 |
| Hypostomus taphorni | GY04173 | MK959902/MK959863/MK959940/MK959990/MK960046 | Guyane, Sawarab bridge | 2 |
| Hypostomus ternetzi | YC164 | JF290462/MK959887/MK959963/MK960016/MK960071 | Argentina, Middle Parana River | 23 |
| Hypostomus uruguayensis | YC10356 | MK959909/MK959886/MK959962/MK960015/MK960070 | Argentina, Middle Parana River | 34 |
| Hypostomus watwata | GF99162 | AJ318352/MK959862/MK959939/MK959989/MK960045 | French Guiana, Oyapok River | 5 |
| Outgroups | | | | |
| Aphanatorulus ammophilus | VZ142 | AJ318346/MK959879/MK959913/MK960006/MK959879 | Venezuela, San Carlos River, near Las Vegas | 2 |

| | | | | |
|---|---|---|---|---|
| Hemiancistrus fuliginosus | UR008 | AJ318359/MK959877/MK959954/MK960004/MK960060 | Brazil, Uruguay River | 3 |
| Pterygoplichthys multiradiatus | VZ119 | AJ318361/MK959878/MK959912/MK960005/MK960061 | Venezuela, Aguaro River | 3 |
| Pterygoplichthys scrophus | 966 | AJ318362/MK959837/MK959910/MK959964/MK960023 | Peru, Loreto, Ucayali, Marañon | 2 |
| Pterygoplichthys zuliaensis | VZ4 | AJ318360/MK959880/MK959911/MK960007/MK960062 | Venezuela, Escalante River | 3 |

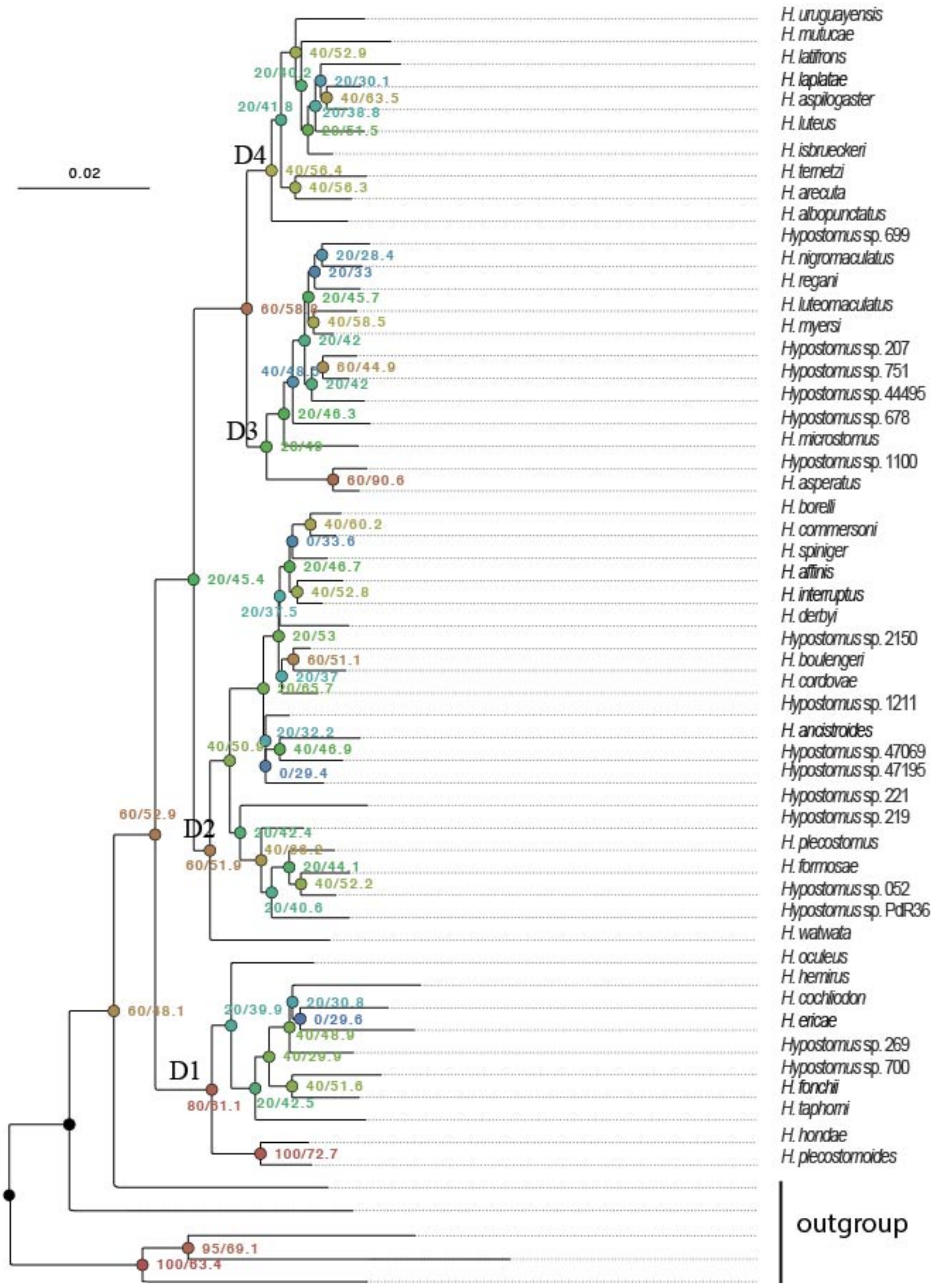


Fig S1. Phylogenetic reconstruction based on the concatenated data showing gCF / sCF in each branch estimated in IQtree. D1 – D4 identify clades.

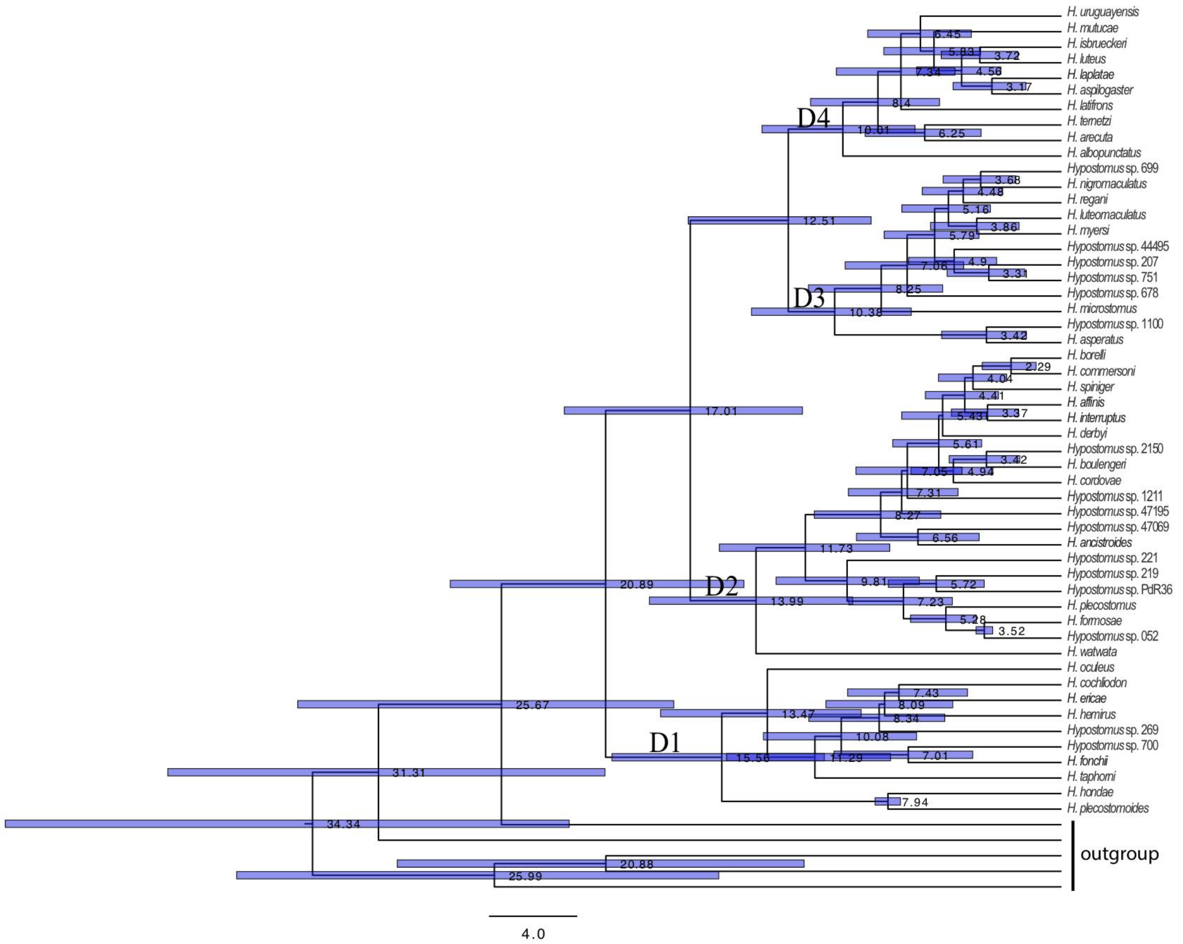


Fig. S2. Time-calibrated multilocus phylogeny for the Hypostomus genus, illustrating mean dates and interval bars (95% HPD) for each node. D1 – D4 identify clades.

Table S2. Models of evolution for the habitat preference in *Hypostomus*. Nr: number of independent transition rates estimated; logLik: the maximum negative log-likelihood; AICc: Akaike information criterion corrected for sample size.

| Model | Nr | logLik | AICc | Δ-AICc |
|---|---|---|---|---|
| ER: equal rates model | 1 | -64.199 | 130.470 | 1.996 |
| SYM: symmetric model | 10 | -51.846 | 128.475 | 0 |
| ARD: all rates different matrix | 20 | -48.074 | 159.481 | 31.006 |

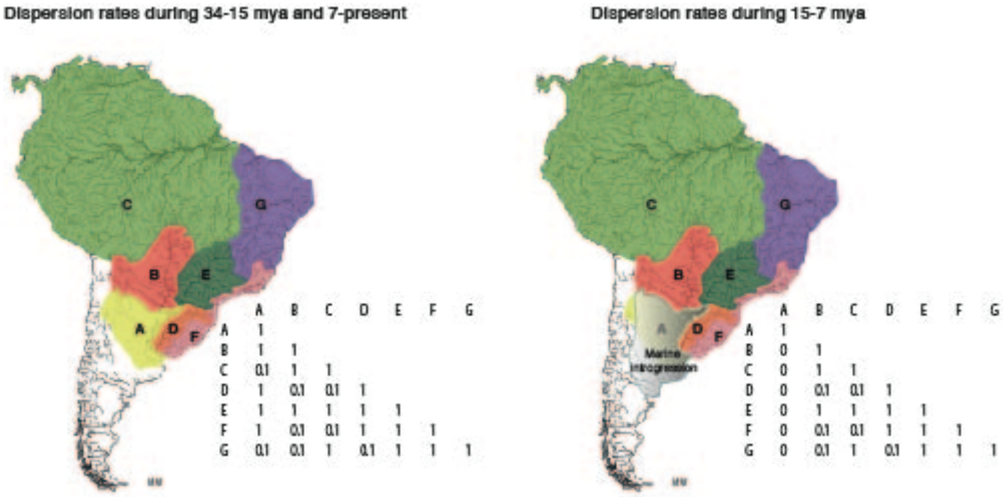


Fig. S3. Dispersal rates between the eco-regions over time used in the DEC analyses The graphical representation of Miocene Marine introgression is approximated .

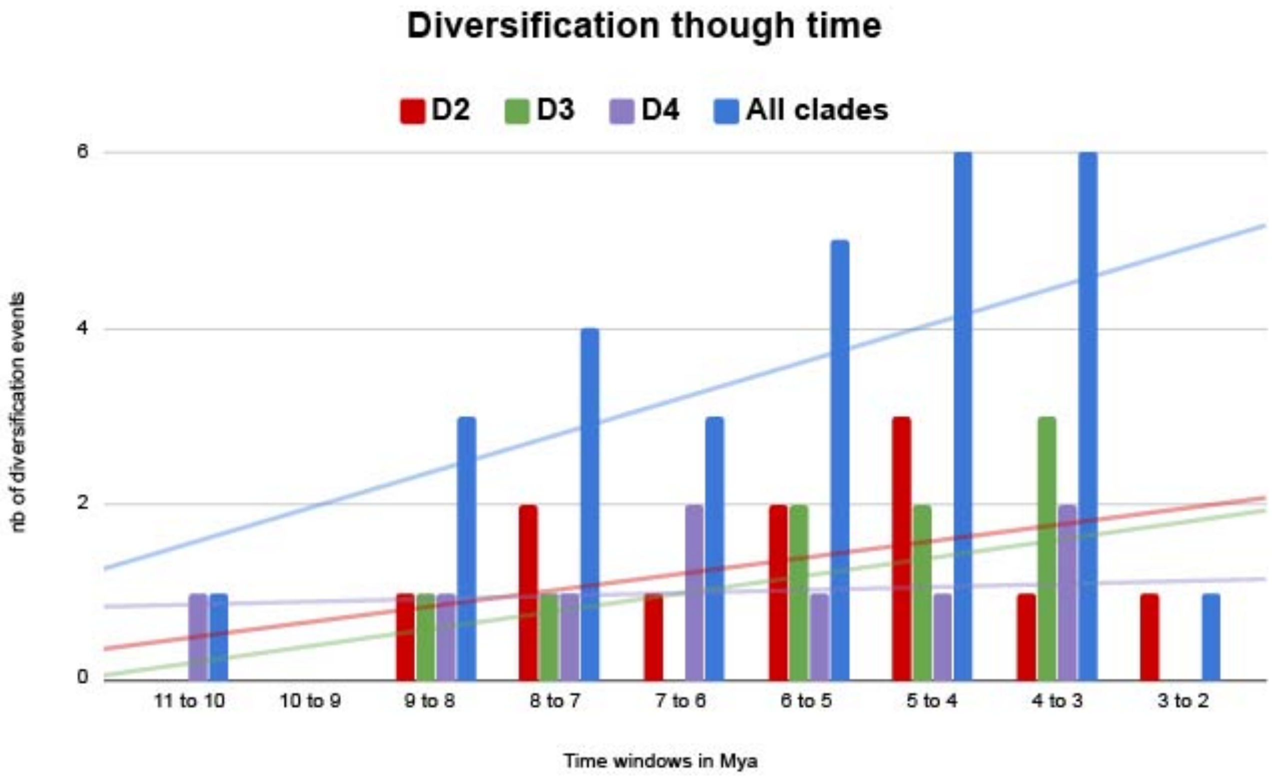


Fig. S4. The number of diversification events for each clade in time windows of one Mya, as well as for the total number of speciation events in *Hypostomus* within La Plata Basin. We also plotted the trend line of each data.
